# The nonlinear structure of linkage disequilibrium

**DOI:** 10.1101/566208

**Authors:** Reginald D. Smith

**Author notes:** Email address (Reginald D. Smith).

## Abstract

The allele frequency dependence of the ranges of all measures of linkage disequilibrium is well-known. The maximum values of commonly used parameters such as *r*^2^ and *D* vary depending on the allele frequencies at each locus. However, though this phenomenon is recognized and accounted for in many studies, the comprehensive mathematical framework underlying the limits of linkage disequilibrium measures at various frequency combinations is often heuristic or empirical. Here, it is demonstrated that underlying this behavior is the fundamental shift between linear and nonlinear dependence in the linkage disequilibrium structure between loci. The proportion of linear and nonlinear dependence can be estimated and it demonstrates how even the same values of *r*^2^ can have different implications for the nature of the overall dependence. One result of this is the value of *D*^*′*^, when defined as only a positive number, has a minimum value of |*r*|. Understanding this dependence is crucial to making correct inferences about the relationships between two loci in linkage disequilibrium.

## 1. Definitions of linkage disequilibrium

Soon after the rediscovery of Mendel’s work and the dawn of modern genetics, one of his laws, his Second Law of the independent assortment of genes, was found to be in conflict with experimental results in several organisms (Bateson Punnett & Saunders (1905, 1906); Morgan (1911)). The concept of linkage, where certain pairs of genes on chromosomes segregated together during recombination much more often than other pairs, became a fundamental concept in theoretical and applied genetics. The mathematical theory of linkage between two loci was first expounded soon afterwards (Jennings (1917); Robbins (1918)). The more general case for multiple loci was derived in detail by Geiringer (1944). The modern formation of the problem, and the definition of the term linkage disequilibrium, was made by Lewontin & Kojima (1960) where *D*, one of the most common measures of linkage disequilibrium, is first defined. Many other measures of linkage disequilibrium would be defined such as *D*^*′*^ (Lewontin (1964)), *r*^2^, the square of the correlation coefficient (Hill & Robertson (1968)), and *d* (Nei & Li (1980)).

Across two bi-allelic loci which have possible alleles *A/a* and *B/b*, the two most well-known measures of linkage equilibrium, are *D* and *r*^2^.

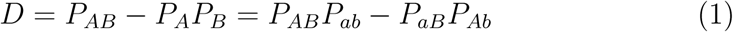

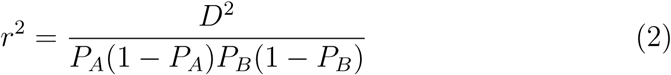

Here *P*_*A*_ and *P*_*B*_ are defined as the frequency of alleles *A* and *B* at each locus and *P*_*AB*_, *P*_*ab*_, *P*_*aB*_, and *P*_*Ab*_ are the probabilities of the haplotypes of the two given alleles at each locus in the same gamete. *D* is equal to the covariance and *P*_*A*_(1 − *P*_*A*_) and *P*_*B*_(1 − *P*_*B*_) are the variances at the respective loci. The range of *D* varies based on loci minor allele frequencies with a maximum of [−0.25, 0.25] where *P*_*A*_ = *P*_*B*_ = 0.5. Given its difficult interpretation, the use of *D* is often modified and represented by the normalized indicator, *D*^*′*^ which normalizes the value of *D* to a range of [0, 1] based on its maximum given the underlying allele frequencies at each locus. For *D* > 0

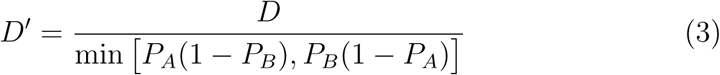

and for *D* < 0

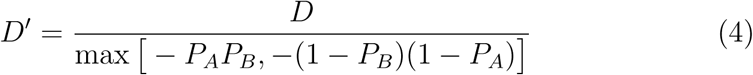

The measures of *r*^2^ and *D*^*′*^ are often used separately or jointly due to the fact they have a common, intuitive, and restricted range of values (Hedrick (1987); Lewontin (1988)). However, there can still be issues of interpretation with both variables. First, though the full range of values can be [0, 1] for each, only *D*^*′*^ can take any value in the range for all values of minor allele frequencies. The ranges of *r*^2^ and *D* are restricted based on the loci allele frequencies. Second, sometimes the two measures can have wildly different, seemingly contradictory, values for the same measurement of linkage disequilibrium. For example, in some cases *r*^2^ can take small values near zero while *D*^*′*^ = 1. This typically happens either when alleles at different loci have a positive correlation but large differences in allele frequencies (e.g. *P*_*A*_=0.91, *P*_*B*_ = 0.06, *D*^*′*^ = 1, *r*^2^ = 0.006) or when alleles at different loci have a negative correlation but similar allele frequencies (e.g. *P*_*A*_=0.91, *P*_*b*_ = 0.94, *D*^*′*^ = 1, *r*^2^ = 0.006; *P*_*b*_ = 1 − *P*_*B*_).

Regarding the first exception, an in-depth analysis on the maximum values of *r*^2^ across different pairs of minor allele frequencies was given by VanLiere & Rosenberg (2008). In the paper, equations 2-5 illustrate the derived maximum values of *r*^2^, dubbed 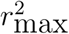, for various minor allele frequency combinations across loci. Depending on the relative values of the major and minor allele frequencies between loci, 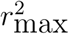 would take the value of either *P*_*A*_(1 − *P*_*B*_)*/P*_*B*_(1 − *P*_*A*_), *P*_*B*_(1 − *P*_*A*_)*/P*_*A*_(1 − *P*_*B*_), *P*_*A*_*P*_*B*_*/*(1 − *P*_*A*_)(1 − *P*_*B*_), or (1 − *P*_*A*_)(1 − *P*_*B*_)*/P*_*A*_*P*_*B*_. One of the key findings of the paper was that 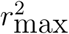 can only be equal to 1, allowing *r*^2^ to take any value, under the two conditions of either *P*_*A*_ = *P*_*B*_ or *P*_*A*_ = 1 − *P*_*B*_. The first condition, *P*_*A*_ = *P*_*B*_, corresponds to positive values of *r*, while *P*_*A*_ = 1−*P*_*B*_ corresponds to negative values of *r*. The description of 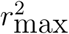 was important for several reasons. For example, it helped give theoretical validation to the long espoused practice of trying to match minor allele frequencies in association studies when inferring a trait locus from a marker locus it is in linkage disequilibrium with. It was known empirically that the limited values of *r*^2^ when minor allele frequencies are mismatched can cause problems with inference and increase the necessary population sizes in association studies (Nei & Li (1980); Kaplan & Weir (1992); Kruglyak (1999); Wray (2005)).

However, the nature of 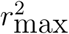 also raises questions about the nature of linkage disequilibrium. Sometimes 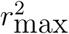 than one. Does this imply that 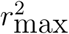 can be extremely limited and much less is the upper bound of any dependence between the two loci or is there additional dependence not encapsulated by *r*^2^? Particularly, should 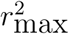 values correspond with complete linkage disequilibrium as defined by *D*^*′*^ = 1 or is the relationship more complicated? In Figure 8 of VanLiere & Rosenberg (2008), there is a clue about a relationship where they demonstrate a proportion of minor allele frequency combinations where 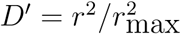 showing that there can often be such a relationship but it is not consistent over all allele frequency values. Where this relation holds and 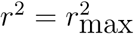 we have complete linkage disequilibrium. However, for those allele frequency combinations where this relationship does not hold, does *D*^*′*^ provide any additional information on dependence beyond that of *r*^2^?

One of the key reasons for the limitations of *r*^2^ and the divergences between *r*^2^ and *D*^*′*^ is that *r*^2^, like all correlation coefficient derived measures, is strictly a measure of the linear dependence between alleles at different loci. On the other hand, *D* and *D*^*′*^ both account for both linear and nonlinear dependence. The issues with *r*^2^ across different allele frequencies is in fact, not just an issue of the definition of *r*^2^ but rather a larger one of nonlinear dependence between loci. This may seem like a subtle or even unimportant detail, however, in this paper it will be demonstrated that the effects of this dichotomy are far from trivial and can demonstrate the error of relying on *r*^2^ alone in certain situations.

## 2. Defining linear linkage disequilibrium

While all values of linkage disequilibrium have a linear component as evidenced by a nonzero value of *r*^2^, how linear the overall relationship is between the two loci should be quantified in order to understand the importance of *r*^2^ in the context of total dependence. As a first step, we will analyze under what conditions the linkage disequilibrium is completely linear and thus has all dependence described by *r*^2^ alone. While the joint probabilities of two alleles at each locus are symmetric, *P*_*AB*_ = *P*_*BA*_ the conditional probabilities are distinct and defined as *P*_*B*|*A*_ = *P*_*AB*_*/P*_*A*_ and *P*_*A*|*B*_ = *P*_*AB*_*/P*_*B*_. Given the conditional probabilities describe the dependence of one allele on the other, we can use them to define the conditions of completely linear dependence.

The linkage disequilibrium between two loci is fully linear when the dual conditional probabilities of any two alleles can be described by a system of linear equations

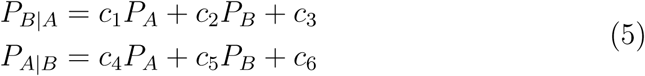

In equation 5 *c*_1_ to *c*_6_ are constants. From the definitions of conditional probabilities and equations 1 and 2 we can derive the two equations

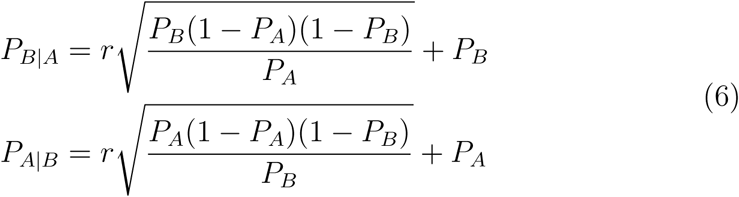

This system of equations is not linear and would seem to imply that there is no case of complete linear dependence, however, there are two conditions where equation 6 reduces to a system of linear equations, in particular the cases of *P*_*A*_ = *P*_*B*_ or *P*_*A*_ = 1 − *P*_*B*_, the identical conditions where 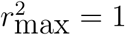. Under both conditions the equations reduce to

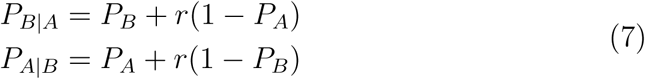

Therefore, only under the conditions of 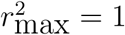 can the linkage disequilibrium be described as completely linear and *r*^2^ captures all dependence. The primacy of *r*^2^ under these conditions of perfectly linear linkage disequilibrium is emphasized by the fact that for these matchings of allele frequencies *D*^*′*2^ = *r*^2^. Further, by Bayes’ Theorem, *P*_*B*|*A*_*P*_*A*_ = *P*_*A*|*B*_*P*_*B*_, we can show that under perfectly linear linkage disequilibrium

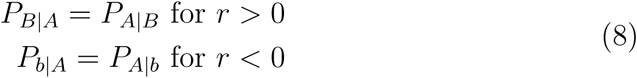

Since the conditions of perfectly linear linkage disequilibrium are extremely narrow, it is now a question of defining the relative strengths of linear and nonlinear dependence in linkage disequilibrium for all other possible combinations of allele frequencies across loci.

## 3. Defining nonlinear linkage disequilibrium

Under the conditions of perfectly linear linkage disequilibrium, *D*^*′*2^ = *r*^2^. However, under all other conditions, *D*^*′*^ can have a range of values for any fixed value of linear linkage disequilibrium as measured by *r*^2^. To understand the ranges of *D*^*′*^ for a given linear contribution from *r*^2^, we can define the minimum and maximum values of *D*^*′*^. For any *r*, the maximum value of *D*^*′*^ is always 1 where linkage disequilibrium measured by *D* is at its maximum value. However, the minimum of *D*^*′*^ can vary. For fixed *r* > 0,

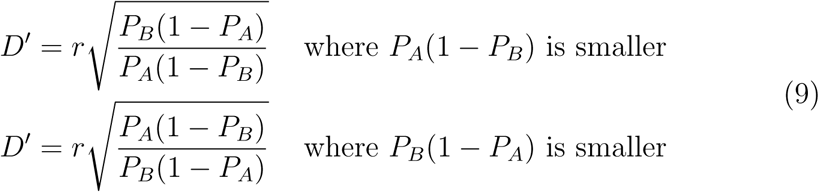

or in the case *r* < 0

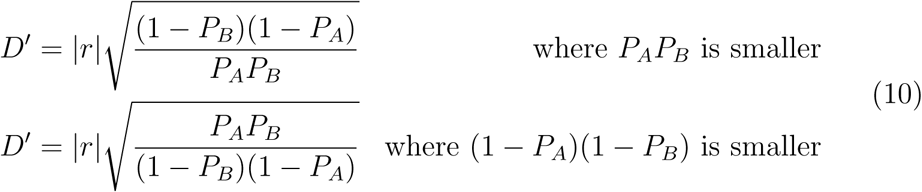

Both possibilities for positive or negative *r* depend on the maximum values of *D* per equations 3 and 4. In both equations 9 and 10, however, the products in the denominator are always smaller than the products in the numerator. Therefore, the square root terms in equations 9 and 10 have a minimum value of one. The maximum of *D*^*′*^ is 1 which means that at complete linkage disequilibrium those square root terms equal 1*/r* and are at a maximum. Therefore, a value of 1 for the square root terms designates the minimum value of *D*^*′*^ giving *D*^*′*^ = |*r*|. Since it is already established previously that *D*^*′*2^ = *r*^2^ designates perfectly linear linkage disequilibrium the following two points can be determined.

First, for a given value of *r*, the range of *D*^*′*^ is limited to [|*r*|,1] where the condition of *D*^*′*^ = |*r*| or *D*^*′2*^ = *r*^2^ indicates perfectly linear dependence between the alleles across both loci where *P*_*B*|*A*_ = *P*_*A*|*B*_ or *P*_*b*|*A*_ = *P*_*A*|*b*_ for positive or negative values of *D* and *r* respectively. Second, for a given value of *r*^2^, any *D*^*′*2^ > *r*^2^ indicates the presence of nonlinear dependence in addition to the linear dependence between alleles across both loci.

Therefore, *D*^*′*2^ more accurately measures full dependence, linear and non-linear. However, the linear and nonlinear proportions of dependence cannot be simply separated just relying on the value of *D*^*′*2^. In order to measure the linear and nonlinear components of *D*^*′*2^ we will rely on the most common and comprehensive measure of total dependence, mutual information. First derived by Claude Shannon when creating information theory (Shannon (1948)) mutual information is a measure of total dependence between two variables and how much reduction in uncertainty we can expect to know about one variable given the other variable. For two bi-allelic loci the mutual information between them is calculated as

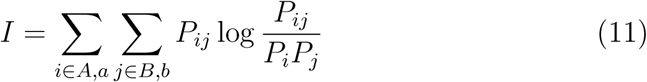

This equation sums the components over all possible allele pairs between the loci. In particular, we can use the mutual information to calculate a ratio of the linear dependence to total dependence, hereafter termed Λ (per Smith (2015)) that will allow us to measure what proportion of total dependence in *D*^*′*^ is linear under any given scenario. For linkage disequilibrium, the mutual information of the base linear case for *D*^*′*2^ can be evaluated by calculating the mutual information for *D*^*′*2^ = *r*^2^. This can then be compared with the total mutual information for any other given value of *D*^*′*2^ > *r*^2^ for the same *r*^2^.

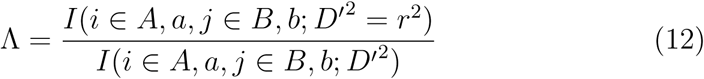

The calculation of both the values of mutual information was done through numerical simulation across different pairs of values for the single allele frequencies *P*_*A*_ and *P*_*B*_. For each pair, the haplotype frequency *P*_*AB*_ and mutual information was calculated for successive different values of *D*^*′*^ ∈ [|*r*|, 1] where *r*^2^ is fixed to calculate the numerator value. Likewise, the denominator was calculated for the same values of *P*_*A*_ and *P*_*B*_ except over all allowable values of 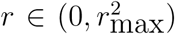. Finally, the ratio indicated in equation 12 is used to calculate Λ.

Based on this procedure we can create heat plots of Λ for different values of *P*_*A*_ and *P*_*B*_. In fact, what we find is regardless the value of *D*^*′*2^, the heat plots are the same with two different plots: one for positive values of *r* and one for negative values of *r*. These are shown in Figure 1.

**Figure 1:**
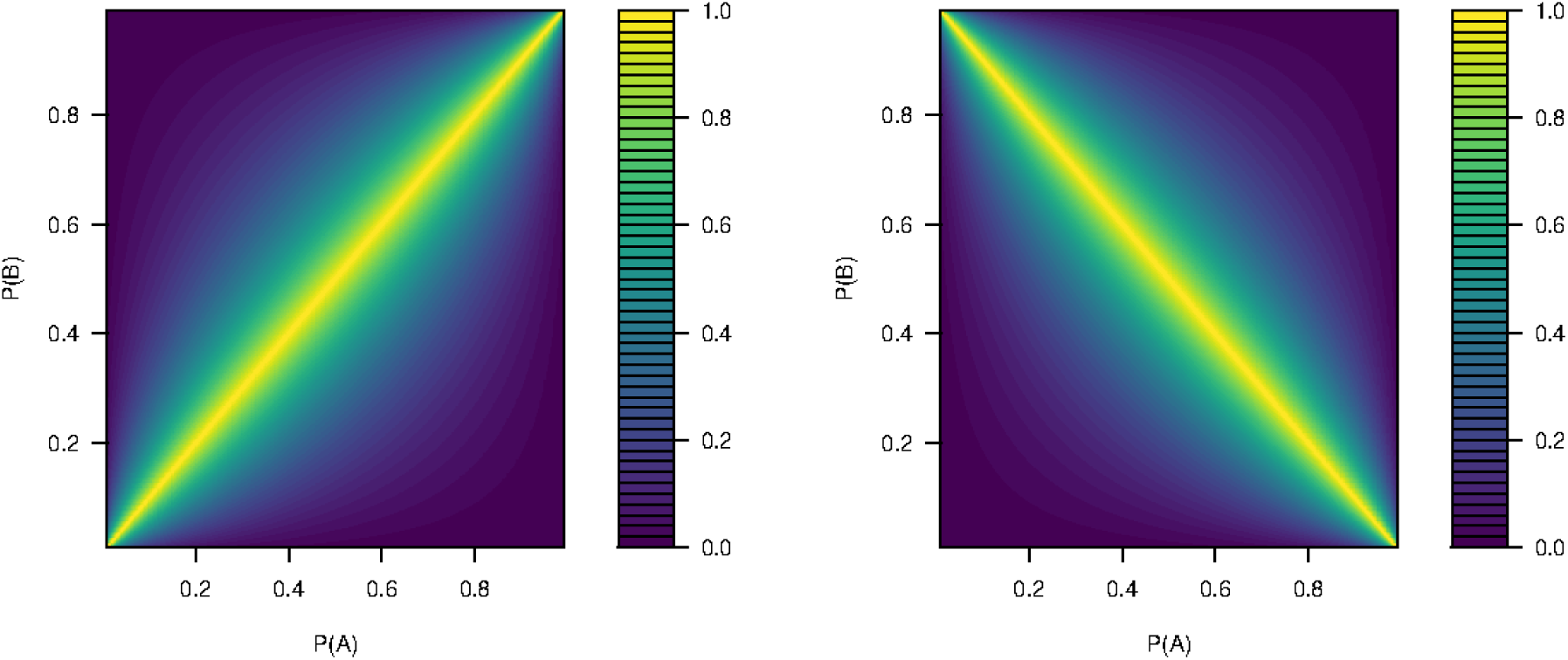
Heat plots of the value of Λ for positive and negative *r* respectively. Yellow indicates a Λ ≈ 1 while violet indicates lower values.

As shown in the plots, for positive *r* the maximum values of Λ, indicating strict linear dependence, only occur in the case of *P*_*A*_ = *P*_*B*_. Outside this region, the linear dependence as a proportion of all dependence rapidly falls off and becomes lower as the ratio of the allele frequencies at each locus become increasingly larger or smaller. Where *r* is negative, we find that the strict linear dependence range changes to where *P*_*A*_ and (1 − *P*_*B*_) are equal. However, like the other situation, the proportion of linear dependence rapidly drops off outside this region.

These plots closely reflect the regions of 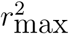 described by Figure 2 in VanLiere & Rosenberg (2008) who plotted the maximum possible value of *r*^2^ by allele frequency pairs for positive or negative correlation. Therefore, we find in a completely different perspective that the proportion of linear dependence to total dependence closely matches the values of 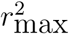 for all allele frequency pairs. Note that Λ keeps the same value for a fixed pair of allele frequencies regardless of the actual value of *r*^2^, even if this value is less than 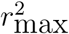.

**Figure 2:**
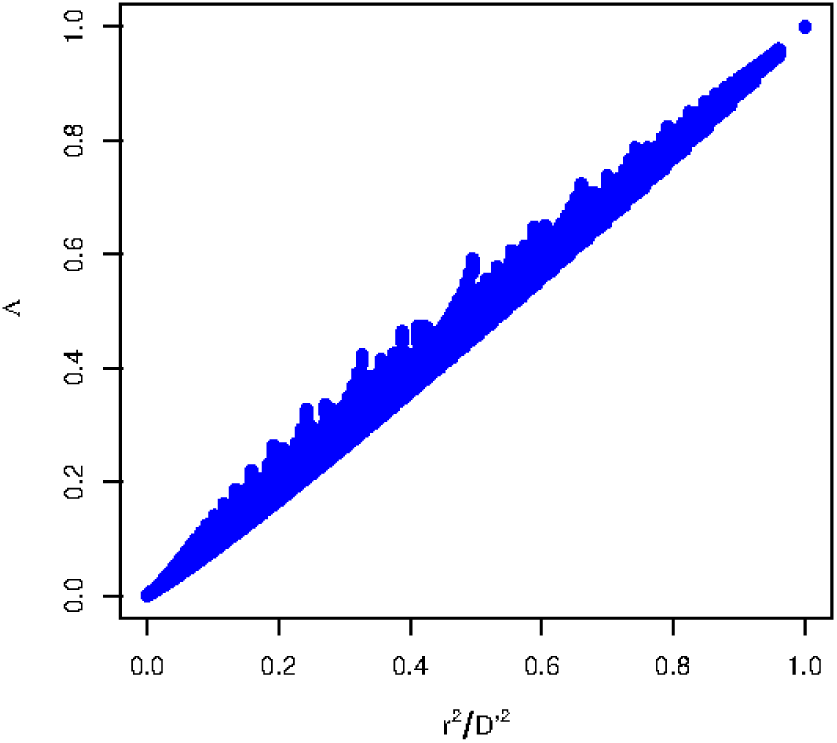
Scatterplot of Λ against *r*^2^*/D*^*′*2^.

## 4. Estimating Λ from allele frequencies

Given the consistent relationships shown in Figure 1 and the fact that *D*^*′*2^ = *r*^2^ for perfect linearity, it is likely Λ is somehow related to *D*^*′*2^ and *r*^2^. What is found is that there is a nearly linear fit across all allele frequencies between Λ and *r*^2^*/D*^*′*2^. This is shown in Figure 2.

Given the definitions of *r*^2^ and *D*^*′*2^, we can show that

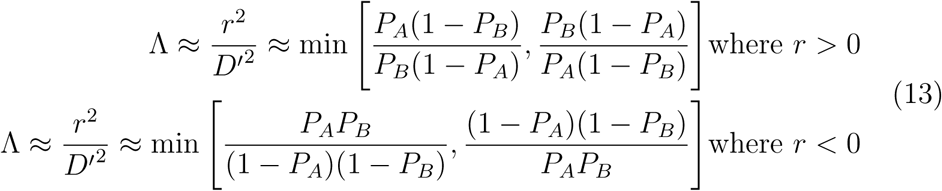

These again match the equations for 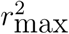 in VanLiere & Rosenberg (2008) across all the values of the allele frequencies. In addition, while in VanLiere & Rosenberg (2008) it was shown that 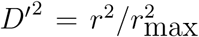 over part of the allele pair frequency pair space, this analysis shows that the value of *D*^*′*2^ ≈ *r*^2^*/*Λ fits across the entire space. Thus, an important finding is that the relative nature of the linear and nonlinear dependence between two associated loci is dictated only by the frequencies of the minor alleles at each locus and not the strength of their linkage disequilibrium. For given allele frequencies, the relative linear and nonlinear proportions of that dependence are identical across all values of linkage disequilibrium.

Having an approximate equation for Λ it is now possible to generalize the relationships between the conditional probabilities *P*_*A*|*B*_ and *P*_*B*|*A*_ for an arbitrary value of Λ. Since *P*_*A*|*B*_ = *P*_*B*|*A*_ or *P*_*A*|*b*_ = *P*_*b*|*A*_ only in perfect linear dependence (Λ = 1), it follows that the introduction of nonlinear dependence fundamentally skews this so that one allele is more dependent on the other for co-occurrence than the reverse. How nonlinear dependence coherently fits into the picture can be shown with Λ. Given equation 13 and Bayes’ Theorem

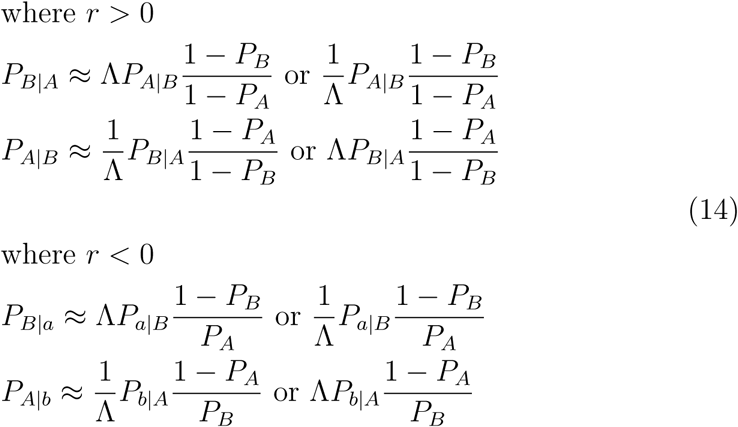

Multiplying this out, regardless of which pair of relations we use, we get the same pair of terms including the joint probabilities.

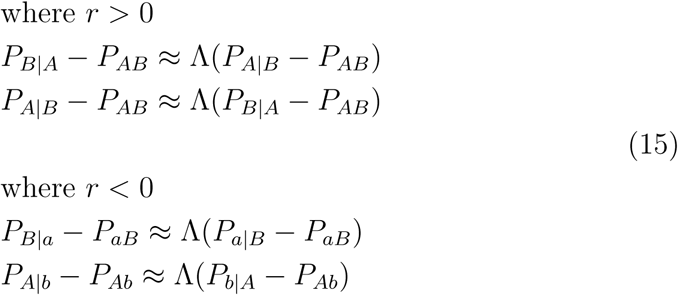

This results in a final consolidated expression decomposing the conditional probability into its linear and nonlinear dependencies

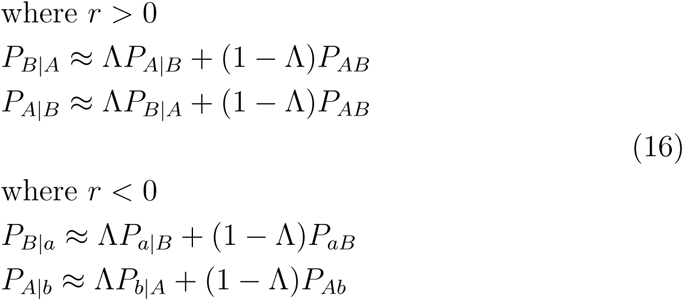

The second term indicates the nonlinear contribution to dependence which gradually reduces the symmetry between co-occurrences at different loci as Λ decreases. At Λ = 0, the dependence is absolute: the conditional probability of finding one allele given the other is equal to their joint probability implying allele one of the alleles has become fixed. Using the first term of equation 16 for further derivation, the equation can also be stated in terms of product instead of a sum

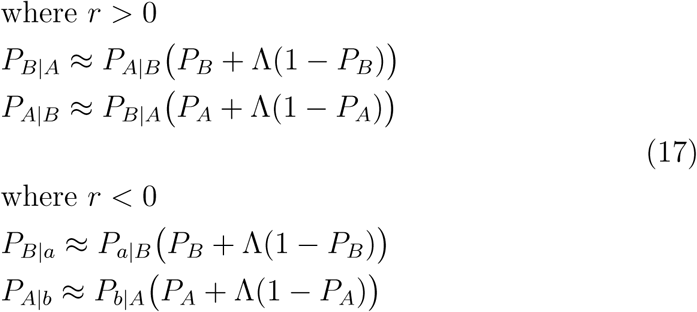

Equations 16 and 17 help illustrate how increasingly nonlinear dependence between alleles at different loci makes using *r*^2^ in calculations problematic where it is explicitly assumed the allele co-occurrences exhibit a symmetric dependence from correlation alone. To illustrate the nature of dependence between alleles at different loci, the diagram in Figure 3 shows the regions of mostly linear, mostly nonlinear, and perfectly linear linkage disequilibrium. Of note is that in the allowed regions of linkage disequilibrium, both the mostly linear and mostly nonlinear regions cover the same area.

**Figure 3:**
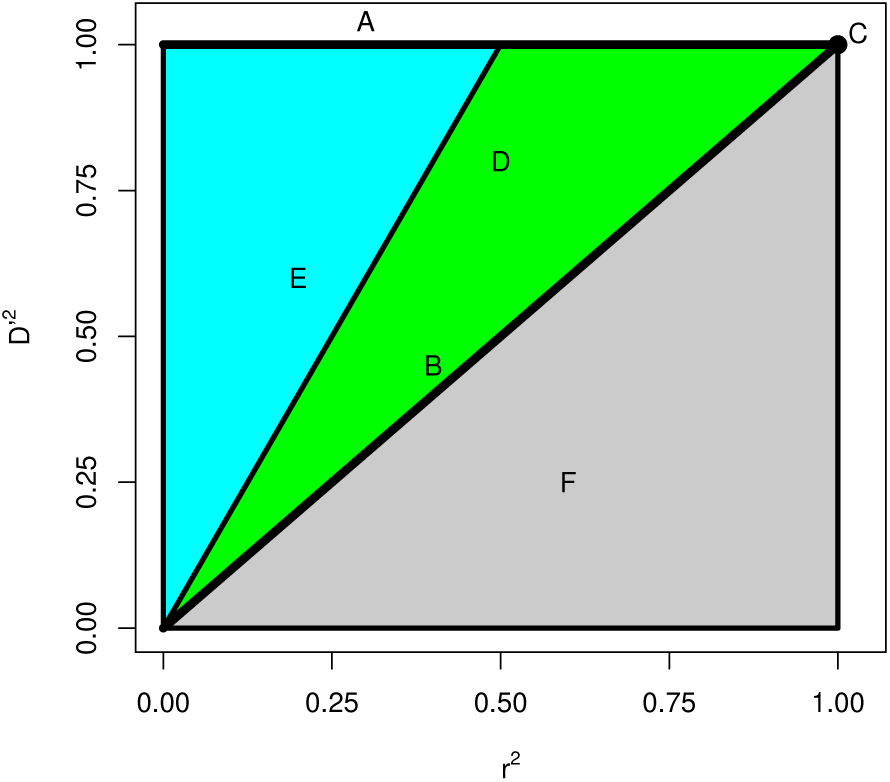
A plot of *D*^*′*2^ vs. *r*^2^ demonstrating the different types of linkage disequilibrium for various values of each measure. Labeled regions indicate (A) complete linkage disequilibrium *D*^*′*^ = 1, (B) perfectly linear linkage disequilibrium *D*^*′*2^ = *r*^2^, (C) perfect linkage disequilibrium *D*^*′*2^ = *r*^2^ = 1, (D) mostly linear linkage disequilibrium (green) Λ ≥ 0.5, (E) mostly nonlinear linkage disequilibrium (light blue) Λ < 0.5, and (F) disallowed combinations of each measure (gray).

In determining the interdependencies of the variances of each locus, the conditional variance between the loci is the key measure. The conditional variance between loci can be calculated similar to the marginal variances as, for example, 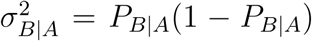 (Dai, Ding, & Wahba (2013)). To demonstrate this using two alleles at separate loci with positive correlation, one can calculate the conditional variance first from *P*_*AB*_ = *D* + *P*_*A*_*P*_*B*_ as *P*_*B*|*A*_ = *D/P*_*A*_ + *P*_*A*_. A subsequent derivation is obtained using the definition of *r* from equation 2.

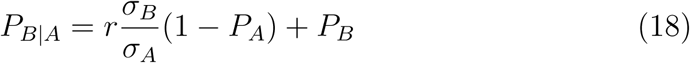

This yields a conditional variance of

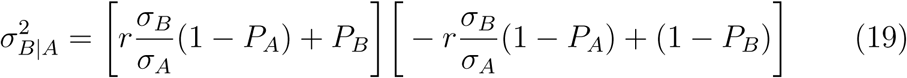

To express the condition variance in terms of Λ, one can use its expression in terms of the site variances.

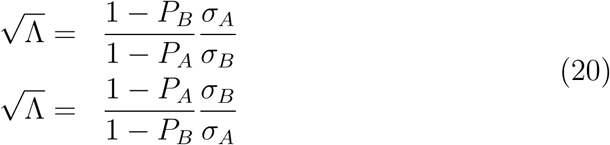

The first and second equations depend on the variance at locus A being relatively larger or smaller than the variance at locus B. The conditional variance is then simplified in terms of the frequency of one locus by one of two expressions.

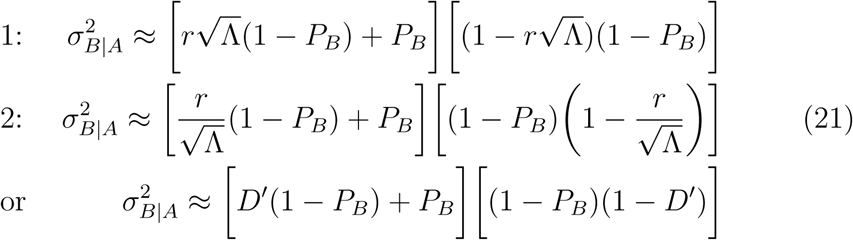

The first and second equation depends on either the variance at locus A or B being relatively smaller. These can be expanded to obtain

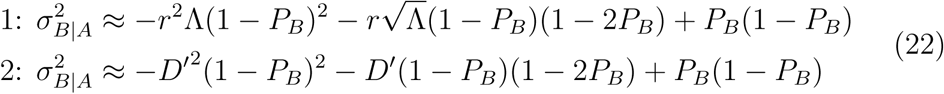

These equations apply for *r* > 0 but when *r* < 0, similar relations apply

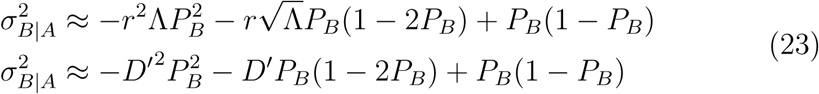

When the relationship is linear with Λ = 1, *P*_*A*_ = *P*_*B*_ = *P*, and 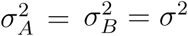 a simplified relationship is obtained

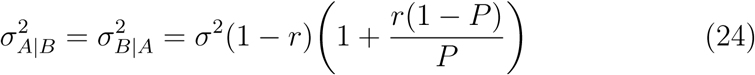

Unlike the canonical case with two conditional normal distributions, the conditional variance does not exactly equal *σ*^2^(1 − *r*^2^) except when *p* = 0.5 and *σ*^2^ = 0.25. However, for most correlations across many allele frequencies the traditional *r*^2^ approximation is still widely valid when the minor allele frequencies at both loci are equal and of moderate frequency. The exceptions will be described in the next section.

## 5. Insights of nonlinear linkage disequilibrium

When the minor allele frequencies at two loci differ, there is nonlinearity in the dependence between the two loci. Therefore, the conditional probabilities of the alleles relative to each other are not symmetric and one allele will give much more information about the presence of the other than vice versa. In extreme cases of weakly linear relationships, this causes a situation where one lower frequency allele at one locus almost always co-occurs with another higher frequency allele at a different locus. This has long been recognized for de novo mutations or rare alleles where *r*^2^ ≈ 0 and *D*^*′*^ = 1 but it can still be applicable at even moderate values of *r*^2^.

The use of *r*^2^ to measure the dependence of the variance between two loci is a proxy for measuring the proportion of each locus’s variance which can be expressed with the conditional variance. In normal regression between two variables with marginal normal distributions, conditional variance is constant across all ranges of both variables and thus a single value, *r*^2^ can characterize all dependence. When this is not the case, as with nonlinear dependence between loci, then the proportion of locus variance that can be explained by conditional variance must be explicitly measured under various conditions. Under any given condition, the proportion of the variance of *B* determined by the variance of *A* can be expressed as

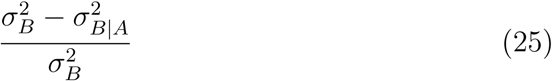

The numerator in the above is the variance of the expected value of *B* given *A*. When the minor allele frequencies are equal at 0.5, this reduces to *r*^2^. When the minor allele frequencies are identical but not equal to 0.5 we have

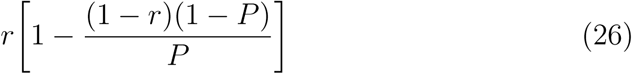

In the above equations, as well as the rest of the derivations in this subsection, *P*_*A*_, *P*_*B*_, and *P* are defined as major allele frequencies. When the allele frequencies differ and nonlinearity comes into play, the value of Λ explicitly defines the proportion of dependence between loci and the relative frequency of each allele at both loci. Where the variance of locus *B* is less than that of locus *A*, increasing nonlinearity means that the presence of *B* almost always signifies the presence of *A*, but *A* only ambivalently matches *B*. The proportion of the variance of *B* that is determined by the variance of *A* increases with increasing nonlinearity so that the proportion is defined as

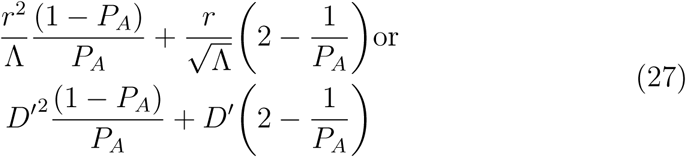

On the other hand, the proportion of the variance of the larger variance locus, *A*, determined by the variance of the smaller variance locus *B*, steadily declines with increasing nonlinearity as the relationship between *A* and *B* is not symmetric. This proportion of variance is described as

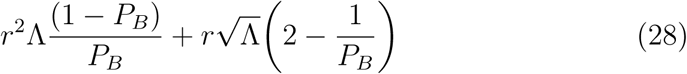

Both these equations represent the proportion of dependence explained when the correlation between the alleles at each loci is positive (*r* > 0).For negative values of *r* the derivation is analogous. Therefore, the values of *r*^2^, *D*^*′*^ and Λ along with allele frequencies can fully characterize the dependence relationships.

In order to visually understand how the relative dependence changes with Λ, Figures 4 and 5 both demonstrate the effects of nonlinearity on loci with unequal variances. The first shows how the overall proportion of variance explained at one locus by the other deviates, higher or lower, from *r*^2^ expectations given the extent of nonlinearity. As demonstrated, the locus with the smaller variance has substantially more of its variance explained by the other locus when linkage disequilibrium is nonlinear.

**Figure 4:**
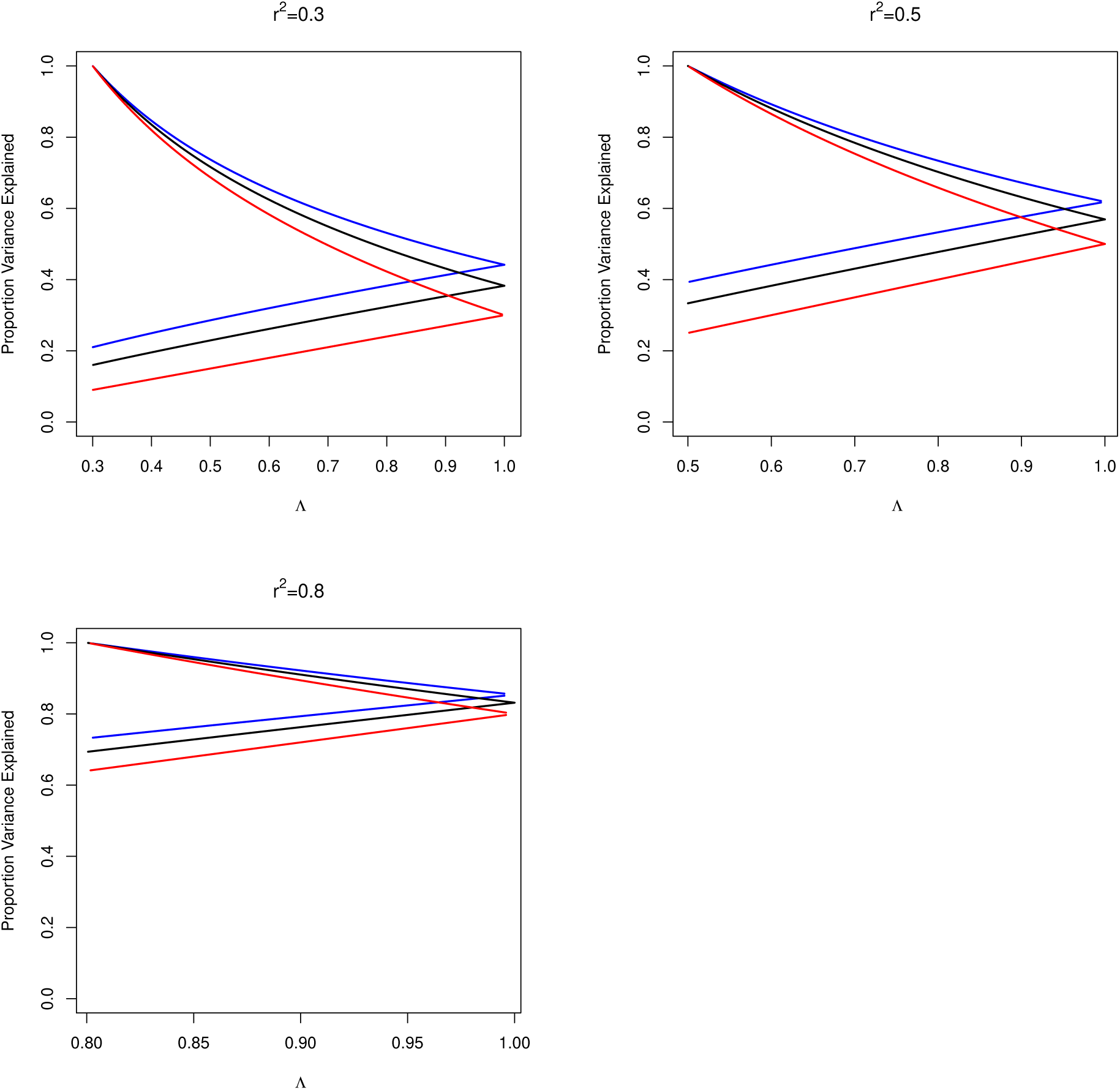
Plots demonstrating the dependence on Λ of the proportion of variance explained at each locus. Each plot represents a stated fixed value of *r*^2^ (*r* > 0) with values of 0.3, 0.5, and 0.8. In each plot are three pairs of lines representing the differing proportions of dependence where the frequency of the major allele at the locus with the largest variance is 0.5 (red), 0.6 (black), or 0.7 (blue). The higher value line of each pair represents the proportion of the variance of the smaller variance locus explained by the larger variance locus. The lower value line in each pair represents the proportion of variance of the larger variance locus explained by the smaller variance locus.

**Figure 5:**
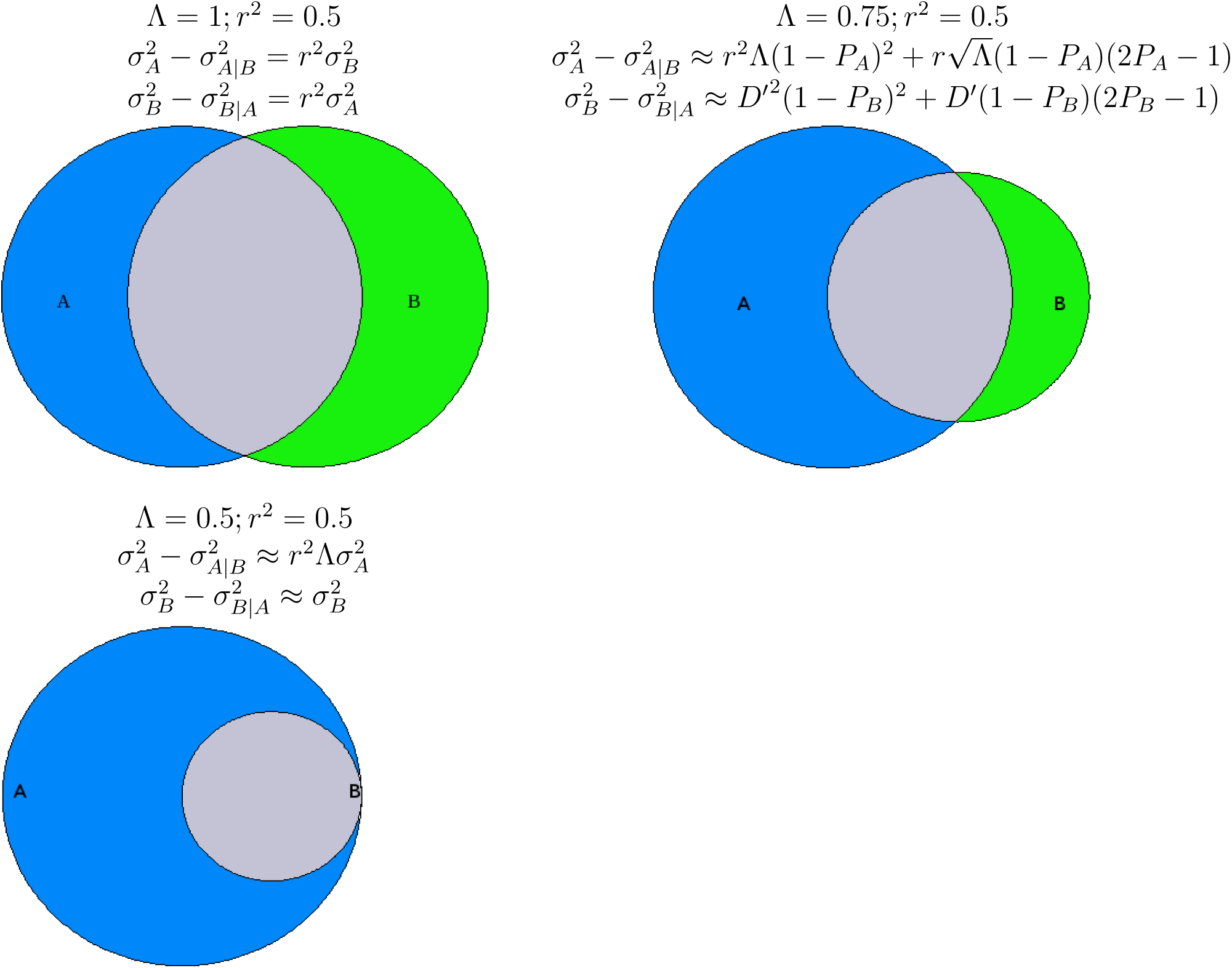
Graphic explanation of how the dependence between loci becomes asymmetric as the linearity, given by Λ < 1, decreases. In all scenarios, *r*^2^ = 0.5 where *r* > 0 at 0.707. Locus *A* retains a major and minor allele frequency of *P*_*A*_ = 0.5 in all scenarios but locus *B* has its major allele frequency increase in order to accommodate the same *r*^2^ for nonlinear linkage disequilibrium. The successive plots show Λ of 1, 0.75, and 0.5. The size of the circles are proportional to the relative conditional variances. The variance at one locus explained by the other is given by the difference between the locus variance and the conditional variance in the equations above the images.

Likewise Figure 5 graphically depicts how much of the variance of each locus is determined by the other by showing how their dependencies overlap. In a manner analogous to a Venn diagram, the amount of overlap for each locus is proportional to the variance explained at that locus by the other. From both loci having the same proportion of dependence explained when Λ = 1 to the point where Λ = *r*^2^ and the smaller variance locus as all of its variance explained by the larger one, the shift in the nature of dependence caused by nonlinearity is clear.

Finally, in Tables 1 and 2 this effect shown numerically for two situations where *P*_*B*_ and *r*^2^ are fixed but Λ is allowed to vary. It is clear that for an increasingly nonlinear linkage disequilibrium, the locus with the largest variance becomes an increasingly less reliable predictor of the other locus with a smaller variance despite all the situations having the same correlation. An additional complication is added when trying to predict how the variance at one locus can be predicted based on the variance at another locus it is in linkage with such as in genome wide association studies.

**Table 1:** The structure of linkage disequilibrium where *r*^2^ = 0.3 (*r* > 0) and *p*_*B*_ = 0.5 for different amounts of nonlinearity.

| $P_A$ | $P_B$ | $r^2$ | $\Lambda$ | $P_{A\vert B}$ | $P_{B\vert A}$ | $\frac{\sigma_A^2 - \sigma_{A\vert B}^2}{\sigma_A^2}$ | $\frac{\sigma_B^2 - \sigma_{B\vert A}^2}{\sigma_B^2}$ |
| --- | --- | --- | --- | --- | --- | --- | --- |
| 0.769 | 0.5 | 0.3 | 0.3 | 1.0 | 0.65 | 1.0 | 0.09 |
| 0.665 | 0.5 | 0.3 | 0.4 | 0.96 | 0.67 | 0.82 | 0.12 |
| 0.665 | 0.5 | 0.3 | 0.5 | 0.93 | 0.69 | 0.69 | 0.15 |
| 0.625 | 0.5 | 0.3 | 0.6 | 0.89 | 0.71 | 0.58 | 0.18 |
| 0.588 | 0.5 | 0.3 | 0.7 | 0.86 | 0.73 | 0.5 | 0.21 |
| 0.555 | 0.5 | 0.3 | 0.8 | 0.83 | 0.75 | 0.42 | 0.24 |
| 0.525 | 0.5 | 0.3 | 0.9 | 0.8 | 0.76 | 0.36 | 0.27 |
| 0.5 | 0.5 | 0.3 | 1 | 0.77 | 0.77 | 0.3 | 0.3 |

**Table 2:** The structure of linkage disequilibrium where *r*^2^ = 0.3 *r* > 0 and *p*_*B*_ = 0.7 for different amounts of nonlinearity.

| $P_A$ | $P_B$ | $r^2$ | $\Lambda$ | $P_{A\vert B}$ | $P_{B\vert A}$ | $\frac{\sigma_A^2 - \sigma_{A\vert B}^2}{\sigma_A^2}$ | $\frac{\sigma_B^2 - \sigma_{B\vert A}^2}{\sigma_B^2}$ |
| --- | --- | --- | --- | --- | --- | --- | --- |
| 0.886 | 0.7 | 0.3 | 0.3 | 1.0 | 0.79 | 1.0 | 0.21 |
| 0.854 | 0.7 | 0.3 | 0.4 | 0.98 | 0.80 | 0.85 | 0.25 |
| 0.824 | 0.7 | 0.3 | 0.5 | 0.96 | 0.82 | 0.74 | 0.29 |
| 0.796 | 0.7 | 0.3 | 0.6 | 0.94 | 0.83 | 0.66 | 0.32 |
| 0.769 | 0.7 | 0.3 | 0.7 | 0.92 | 0.84 | 0.59 | 0.35 |
| 0.744 | 0.7 | 0.3 | 0.8 | 0.90 | 0.85 | 0.53 | 0.38 |
| 0.722 | 0.7 | 0.3 | 0.9 | 0.88 | 0.86 | 0.48 | 0.41 |
| 0.7 | 0.7 | 0.3 | 1 | 0.86 | 0.86 | 0.44 | 0.44 |

### 5.1. Inference in association studies using high frequency alleles

It is standard process in many analyses such as association studies to measure the variance of groups of single nucleotide polymorphisms (SNPs) with respect to a trait being studied. The SNP density along chromosomes is often chosen with the understanding that even if the SNPs themselves are not causal sites, the causal sites, often termed quantitative trait loci (QTL), are likely in linkage disequilibrium with multiple SNPs and thus SNP variance with respect to trait variance is related to the causal site variance with respect to the trait. The typical expectation is that given a casual site’s variance, 2*p*(1 − *p*), the SNP variance with respect to the variance of the QTL is 2*p*(1 − *p*)*r*^2^.

However, as demonstrated in the previous section, the conditional variance at one locus given the other is not always straightforward given *r*^2^ alone. In fact the variance of the SNP with respect to the variance of the QTL should only be 2*p*(1 − *p*)*r*^2^ under the narrow condition of the minor allele frequency at both loci being 0.5. Assuming the locus of the SNP has a larger variance than that of the putative QTL, often the case especially if the QTL is expected to have low frequency or rare (minor allele frequency ≤ 1%) alleles, nonlinearity should be assumed to be present to some extent in most analyses.

Following on the notation and analysis of Weir (2008) for a quantitative trait under investigation, *Y*, the SNP marker locus will be designated *M* and the causal site/QTL will be designated *T*. The mean value of *Y, µ*_*G*_, is defined using the genotype scores *G*_*TT*_, *G*_*Tt*_, and *G*_*tt*_.

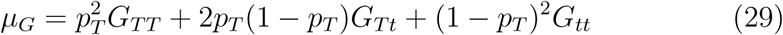

The additive and dominance variances of *T* are similarly defined as

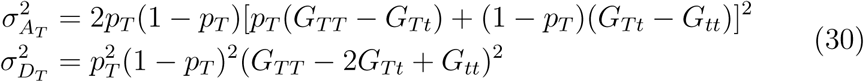

The variance of *Y* is 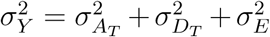 where 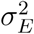 is the environmental variance. The expected conditional trait means for *Y* given each marker genotype given a marker-trait correlation of *r* are

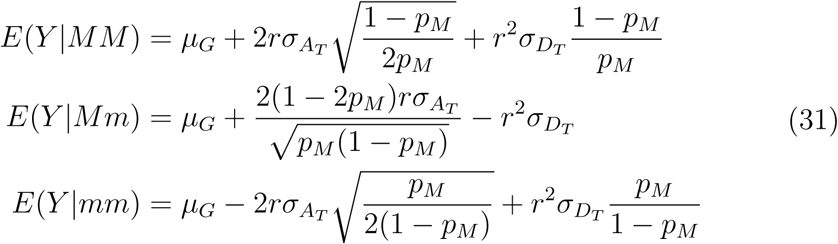

In order to investigate the conditional variance of *Y* given the presence of marker allele *M*, we must derive the expected mean of *E*(*Y* |*M*). First we calculate the expected joint probability *E*(*Y M*) and then *E*(*Y* |*M*).

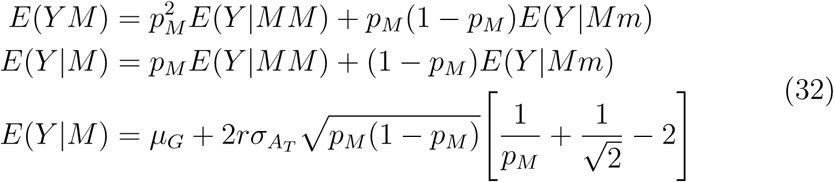

The dominance variance terms cancel when looking only at the conditional variance of one of the marker alleles. We can thus derive the conditional variance 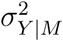 from the expected mean.

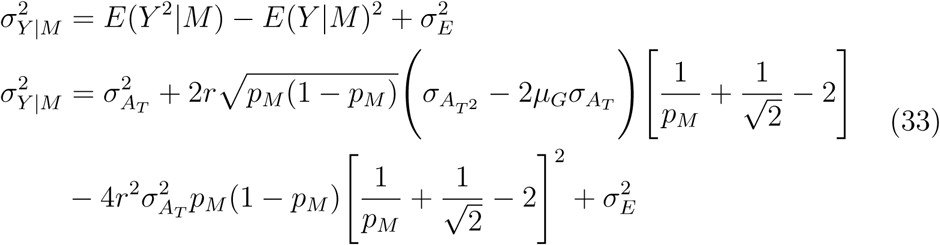

When *p*_*M*_ = *p*_*T*_ = 1*/*2, the term 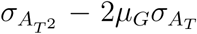 (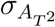 is the additive variance of *Y* ^2^) simplifies to 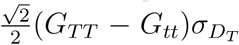. If dominance variance is assumed to be absent and the trait is completely additive we have at complete linearity

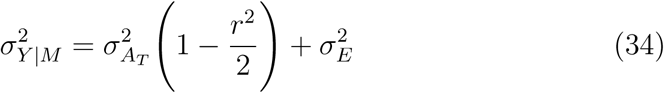

For the general case of 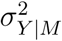, under the condition of *r* > 0 and 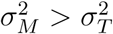, we can incorporate Λ as

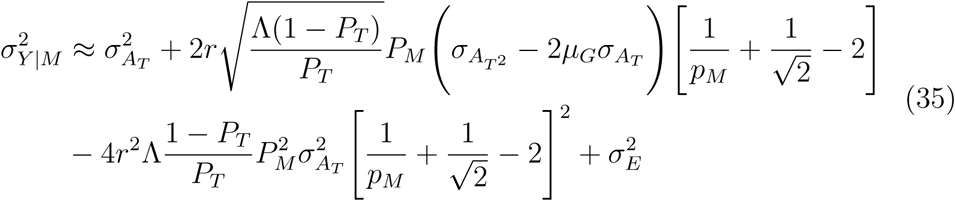

Similar to the association between each loci, the amount of variance of *Y* explained by the marker variable is more complex when the marker and trait loci variances are unequal again becoming only a simple dependence on *r*^2^ only when *P*_*M*_ = *P*_*T*_ = 1*/*2.

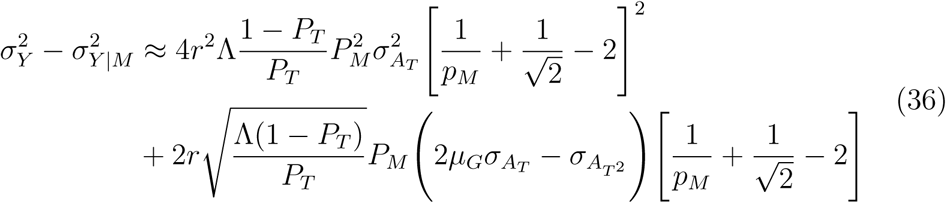

While the above can be accounted for by using both the marker and trait loci variances, often the trait loci are unknown and dependence is inferred based off the correlation between the trait values and the marker. However, while it is recognized that weak linkage disequilibrium and a small variance in a putative trait locus reduce the value of the *r*^2^, the same value of *r*^2^ can still deliver widely different types of dependence depending on the value of Λ.

A final note regards the regression of *Y* against a scored variable of the marker locus *X* where similar to *Y*, the mean and additive variance are defined as

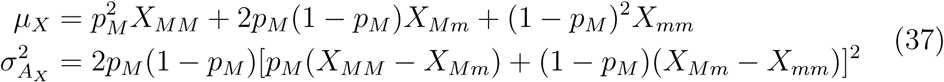

The expected conditional mean and the conditional variance of *Y* given *X*, where *r*_*MT*_ is defined as the correlation between the marker and trait loci, are defined as

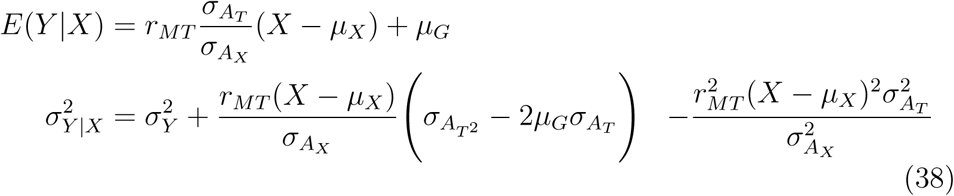

When linearity conditions are met and 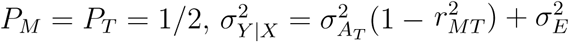. Finally, one can also define the proportion of the variance of *Y* explained by *X* following

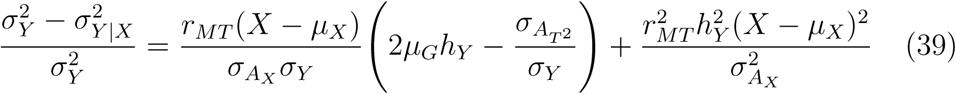

The variable 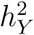 is the narrow heritability of *Y* in the population and environmental conditions under study. Under linear conditions at *P*_*M*_ = *P*_*T*_ = 1*/*2, the proportion of variance of *Y* explained by *X* is 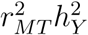. While the correlation between *X* and *Y* is still accurately estimated as *r*_*MT*_ *h*_*Y*_, like the example of two loci, the overall dependence between *X* and *Y* is not exactly the simple relationship of 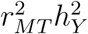 except under narrow conditions.

## 6. Nonlinearity of linkage disequilibrium under evolution

Besides an additional descriptor of the dependence structure between a pair of loci, the relative linearity of linkage disequilibrium can show distinct patterns under some conditions of evolution. In addition, because it is based only on allele frequencies, Λ is more resistant to the decay effects of recombination compared to linkage disequilibrium measures. *D*^*′*2^ and *r*^2^ decay due to recombination at the same rates while the effects of recombination on Λ generally cancel out except for selective or stochastic effects on allele frequencies.

### 6.1. Selective sweeps and hitchhiked alleles

A common scenario implicated in the formation of linkage disequilibrium between pairs of neutral alleles is their hitchhiking with a third allele they are in linkage with whose frequency is increased by a (hard) selective sweep (McVean (2007); Jones & Wakeley (2008); Pfaffelhuber, Lehnert, & Stephan (2008)). Under this scenario, patterns of linkage disequilibrium on either side of the locus undergoing selection tend to show increasingly higher levels until the beneficial allele reaches fixation. Afterwards, recombination begins the inexorable decay of the correlation between the loci. The nonlinearity of linkage disequilibrium does not directly affect nor modify this process, however, barring evolutionary forces that change the neutral allele frequency after the fixation of the selected allele, Λ remains relatively constant until the linkage disequilibrium disappears entirely and it becomes essentially meaningless. It can be demonstrated that despite the various values of the linkage disequilibrium in neighboring sites generated by the sweep, the linearity increases, often from nearly nonlinear to nearly linear, and that this pattern of linearity is preserved in the linkage disequilibrium even after fixation or the relaxation of selection and recombination begins to degrade *D*^*′*^ and *r*^2^.

The relative resilience of Λ is shown in Figure 6 for two pairs of neutral alleles with different recombination probabilities on the same side of the selected site. A selective sweep with adjacent loci at various distances based on recombination frequency *c* was simulated in Python with SimuPop (Peng & Kimmel (2005)). Both pairs of alleles were included in the same simulation run. The strong positive selection on the selected allele leads to fixation around generation 60 where linkage disequilibrium peaks and *r*^2^ and *D*^*′*^ begin to decay. Λ shows no such change, however, given the balance of forces maintaining allele frequencies after fixation. Therefore, many generations after the selective sweep, Λ is a marker of the character of linkage disequilibrium at the time of fixation.

**Figure 6:**
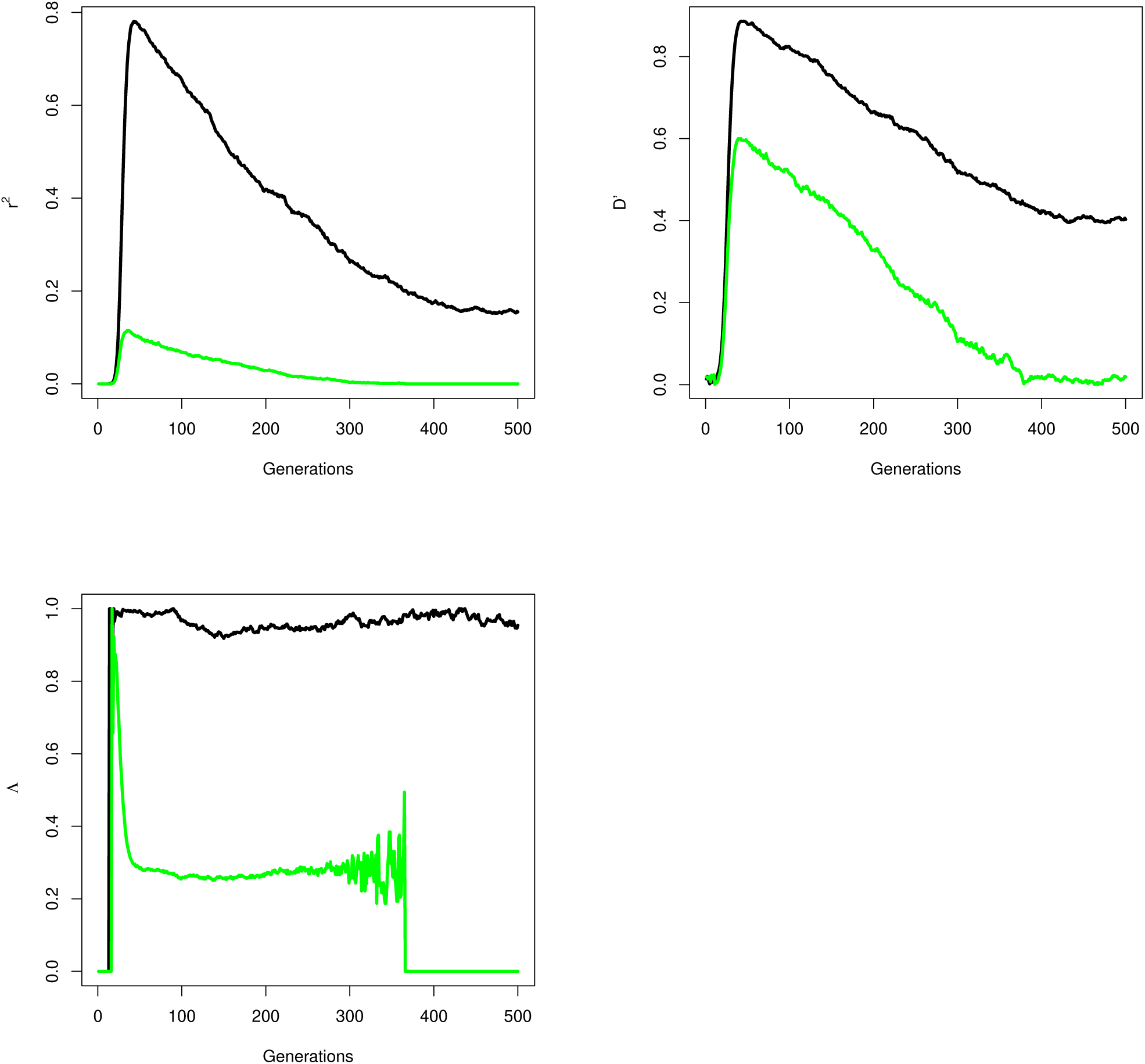
Graphs over 500 generations of the linkage disequilibrium measured by *r*^2^ and *D*^*′*^ as well as Λ for two pairs of neutral alleles hitchhiking an allele under strong selection that becomes fixed in 59 generations. The pairs have recombination frequencies *r* = 0.002 (black lines) and *r* = 0.004 (green lines). Population is *N* = 100, 000 with selection *s*=0.8.

In a region where a selective sweep is expected to have caused the fixation of an allele in the past, a signature of the former association could be the level of linearity in linkage disequilibrium in surrounding regions, assuming the relative stability of the neutral allele frequencies since the sweep. In tandem with searching for linkage disequilibrium, which may have substantially decayed, regions of neutral alleles that are located relatively closely in the chromosome and share relatively similar minor allele frequencies can represent a distant echo of a past sweep.

Assuming the association was once strong, where *D*^*′*^ = 1, a rough measure of the number of generations since the beneficial mutation was fixed could be given by

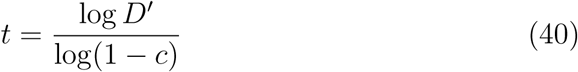

An equivalent expression using the *r* instead is

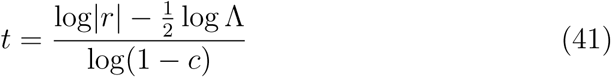

These assumptions are similar to current approximations used to time the decay of linkage disequilibrium. However, understanding the dynamics of Λ allows a different estimation for cases when linkage disequilibrium has substantially decayed and is extremely low. After the sweep, linkage disequilibrium will slowly decay until it reaches the level of linkage disequilibrium expected from drift-recombination equilibrium. This was first discovered through calculation in Hill & Robertson (1968) as *r*^2^ ≈ 1*/*4*N*_*e*_*c* as *N*_*e*_*c* increases. As this is the background linkage disequilibrium expected between any two loci, substituting 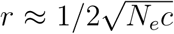 in equation 41 gives us a minimum time since the fixation of the swept allele as

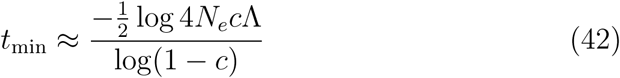

Therefore, even at minimal levels of linkage disequilibrium, one can use the behavior of Λ and assumptions about linkage disequilibrium at the time of the fixation of the sweep to estimate the minimal time elapsed since fixation. Of course this equation assumes the sweep occurred long enough ago to allow the linkage disequilibrium to be decayed by recombination but not so long ago that neutral mutations have begun to significantly alter the allele frequencies at either of the pair of neutral loci. It also requires a reasonable estimate, or range of estimates, for the effective population size, which is assumed to be constant.

## 7. Conclusion

Linkage disequilibrium is a useful and ubiquitous aspect of all genomes. While its measurement given frequency data may seem relatively straightforward, it would behoove all researchers to understand that the measurement of a value of linkage disequilibrium with any preferred measure may mask underlying complexities in the relationship between the loci in question. One measure, be it *r*^2^ or *D*^*′*^, cannot reconcile these issues. In particular, the realization that *r*^2^ measures only linear dependence and is not the same for different allele frequency pairs is essential. Theoretical and experimental analyses based on assumptions of only linear dependence may inadvertently skew estimates of frequency or variance dependence between loci pairs.

Given the crucial assumptions regarding linkage disequilibrium in association studies, it is important to make realistic calculations as to how strong the linkage disequilibrium is between the marker locus and an unidentified causal trait locus given the variance of a marker in relation to the variance of the trait being studied. The key point is while it is well recognized that mismatches in variance between loci reduce the possible strength of linkage disequilibrium as measured by *r*^2^ it is not accurate to view all identical values of *r*^2^ across loci pairs of different respective variances as measuring dependence in the same manner and thus caution is warranted in making inferences on the amount of variation the underlying trait locus is actually inducing on the marker locus in what seem to be similar linkage disequilibrium conditions. Especially for marker loci with relatively high minor allele frequencies, linkage disequilibrium with trait loci with small variances due to rare alleles will likely have a conditional variance, and thus dependence, less than *r*^2^ alone dictates.

Given it is calculated based on allele frequencies, Λ does not have any known direct evolutionary import though it can serve as a marker for past evolutionary events at neutral loci where recombination alters the structure of the main linkage disequilibrium measures. It can also possibly serve as another informative measure of the difference in allele frequencies between loci.

Understanding linkage disequilibrium is a crucial concept required to understand a wide variety of effects from evolutionary forces as well as the underlying population structure and genomic architecture. Hopefully understanding its nuances across all possible ranges of effect can allow it to become even more useful and accurate in the future.

Declarations of interest: none. This research did not receive any specific grant from funding agencies in the public, commercial, or not-for-profit sectors.

